# Health Information Needs and Health Seeking Behavior during the 2014-2016 Ebola Outbreak: A Twitter Content Analysis

**DOI:** 10.1101/103515

**Authors:** Michelle L. Odlum, Sunmoo Yoon

## Abstract

**Introduction:** For effective public communication during major disease outbreaks like the 2014-2016 Ebola epidemic, health information needs of the population must be adequately assessed. Through content analysis of social media data, like tweets, public health information needs can be effectively assessed and in turn provide appropriate health information to effectively address such needs. The aim of the current study was to assess health information needs about Ebola, at distinct epidemic time points, through longitudinal tracking.

**Methods:** Natural language processing was applied to explore public response to Ebola over time from the beginning of the outbreak (July 2014) to six month post outbreak (March 2015). A total 155,647 tweets (unique 68,736, retweet 86,911) mentioning Ebola were analyzed and visualized with infographics.

**Results:** Public fear, frustration, and health information seeking regarding Ebola-related global priorities were observed across time. Our longitudinal content analysis revealed that due to ongoing health information deficiencies, resulting in fear and frustration, social media was at times an impediment and not a vehicle to support health information needs.

**Discussion:** Content analysis of tweets effectively assessed Ebola information needs. Our study also demonstrates the use of Twitter as a method for capturing real-time data to assess ongoing information needs, fear, and frustration over time.

All authors have seen and approved the manuscript.

## Introduction

The 2014-2016 West African Ebola outbreak was the most pervasive in history, reaching epidemic proportions^1^. The outbreak was marked with 9,729 fatalities, 23,948 reported cases and uncontrolled spread ^2^. It was further characterized as an epidemic of fear and anxiety, when case identification reached the US and Europe ^3^. In spite of reports describing modes of transmission, signs and symptoms and the low likelihood of a widespread epidemic, people in developed nations remained overly anxious. In fact, Americans listed Ebola as the third top health concern in October of 2014, clearly indicating fear and highlighting the need for effective health education^4^. The Conceptual Framework of Public Health Surveillance and Action (PHSA) indicate that public health surveillance and action to control disease are connected through data information messages^5^. To support public health action, health information messages must be widely disseminated to the public. As fear drives epidemics, evidenced by the Ebola spread in West Africa, it is essential to understand public opinion to identify health information needs. When these needs are recognized, tailored literacy appropriate health information can be disseminated to diminish widespread fear^6^. Social media can support dissemination efforts. It allows tremendous opportunities to provide literacy appropriate health information through mass dissemination^8^. Social Networking Sites like Twitter encourages users to provide and share information. The analysis of social media data allows for an assessment of the public’s knowledge, personal experiences, and health information needs^7^.

Mining of social media data provides a snap shot of health knowledge in addition to the longitudinal tracking of changes in such knowledge and emerging health information needs over time. Micro-blogging is a very powerful and popular communication tool. Millions of messages are sent daily on micro-blogging sites such as Twitter^8^. Research has demonstrated that Twitter is a reliable source for tracking knowledge and opinions for a variety of events and issues^7^. Such tools allow the public to play a role in knowledge translation including information generation, filtering and dissemination^7, 9^. It then becomes critical during epidemics and other emergencies to monitor online public perceptions and responses. Online monitoring allows for the examination of knowledge translation and for the modification and tailoring of health information for effective health educational campaigns^5, 7^.

The current study sought to assess health information needs about Ebola through longitudinal tracking. We provide a snapshot of beliefs, opinions and responses at three time points beginning in August 2014, when the outbreak raged out of control and through March 2015, when the outbreak showed evidence of containment. Through natural language processing (NLP) and content analysis, we explore how Twitter users communicate about Ebola-related events over time and identify health information needs that emerge at each time point.

## Methods

### Tweet Corpus

Tweets mentioning Ebola were collected longitudinally from Twitter (https://twitter.com/https://twitter.com/) via Google Chrome-based version of NCapture^TM^ a crawler that captures internet-based text including social media for the analysis of data in NVivo. The direct access of tweets (tweet text) through https://twitter.com provides the streaming Application Programing Interface (API)^19^. Streaming API allows its users to capture only limited amount of all tweets (e.g., 18000 Tweets per 15 minutes), the domain experts (SY, MO) captured tweets daily for optimal access to available tweets from July 2014 to March 2015.

To assess the change in Ebola information needs, we analyzed tweets one week after the official CDC announcement of the Ebola outbreak in West Africa. We additionally analyzed tweets three months later after increased efforts to fight the epidemic and increase public awareness of the virus. We also analyzed tweets 6 months after the CDC announcement, when new cases were on a steady decline. At this time point, Liberia was reporting no new cases for two consecutive weeks. In Conakry, Guinea and Freetown Sierra Leone, the outbreak formed geographic contagious arcs enabling response efforts focused on smaller areas^17^. Specifically, the three time points categorized by the domain experts 1) baseline: one week after the official July 28^th^, 2014 CDC announcement of Ebola (August 4^th^ to 8^th^, 2014); 2) 3-month post: three months after the official CDC announcement of Ebola (October 23^rd^ to 26^th^, 2014) and 3) 6-month post: six months after the official CDC announcement of Ebola (March 3^rd^ to 7^th^, 2015).

Using Ebola as the keyword, we retrieved all available English and Spanish language tweets. All Ebola-related hashtags were daily monitored for its mutation by the domain experts and any mutated keywords were included in the data analysis. Examples of included hashtags are #Ebola, #EbolaOutbreak, #EbolaVirus and #EbolaFacts. All Ebola-related hashtags (both simple and complex) were considered. Tweets were included in the analysis if they referred to Ebola. Tweets were collected once to every 15 minutes for 4-5 days for the three identified time points based on the total number of the tweets. This allowed for the collection of a representative sample to overcome the challenges of time and amount limit (e.g., 18000 tweets per every 15 minutes). For example, we did not need to extract tweets 10 times per day when Ebola was rarely mentioned. On the other hand, when Ebola tweets spiked in frequency, 15 minute data collection was required by the domain experts. Data elements collected for each tweet included content, time stamps, geographical locations and self-identified locations, the user names, the message type (unique or retweet), the number of Twitter followers. Dissemination is defined as the number of tweets and subsequent retweets spread to Twitter followers.

**Natural language processing** was conducted to depict the topics discussed about Ebola. To identify topics of collected tweets, we cleaned symbols and web addresses, and transformed text to a vector form and N-gram. This followed by reducing the dimensionality of the volume using Notepad++ and Weka 3.7. Tweet cleaning included the removal of nonstandard or special characters that may hinder the use of content mining tools, including the removal of hashtags. Transferred text is represented as a vector of features. N-grams, an example of features, is a subsequence of N items in a given sequence from characters to words. A tweet term-frequency dictionary was computed with the N-gram method. To reduce the dimensionality to algorithmically process the data, stop words (e.g., and, to, for) were removed using the stop-word removal function in Weka. Stemming, the process of identifying a word root and removing suffixes and prefixes was also performed to remove dimensionality. The stemming algorithm Porter’s algorithm, an affix-removal approach, was applied through “snowball stemmers” within Weka^10^. Preprocessed data was then analyzed to discover patterns including hot topics and sentiment. Topics were detected and summarized through descriptive statistics (e.g., frequency count of specific tweets), visualization, classification and clustering. The N-gram forms (unigram, bigram and trigram) of tweet messages were clustered based on content similarities for topic detection. K-means algorithm was then applied using Weka. Clusters were visualized to summarize the detected topics using infographics. Tweets were also thematically categorized based on their content and user information. Users were further coded based on world regions (i.e., Africa, Asia, Europe, Middle East, North America and South America).

### Results

#### Information Spread

Time point 1: After the first official CDC announcement telebriefing, 45,079 tweets were posted and disseminated to 11,698,549,157 Twitter followers from August 4^th^ to 8^th^, 2014. Time point 2: From October 23^th^ to 26^th^, 2014, 3 months after the first announcement a total of 66,552 tweets were posted and disseminated to 31,265,701,152 Twitter followers. Time point 3: From March 3^rd^ to 7^th^, 2015, 6 months after the first announcement a total of 44,016 tweets were posted and disseminated to 11,141,339,478 Twitter followers (Figure 1).

**Figure 1.**
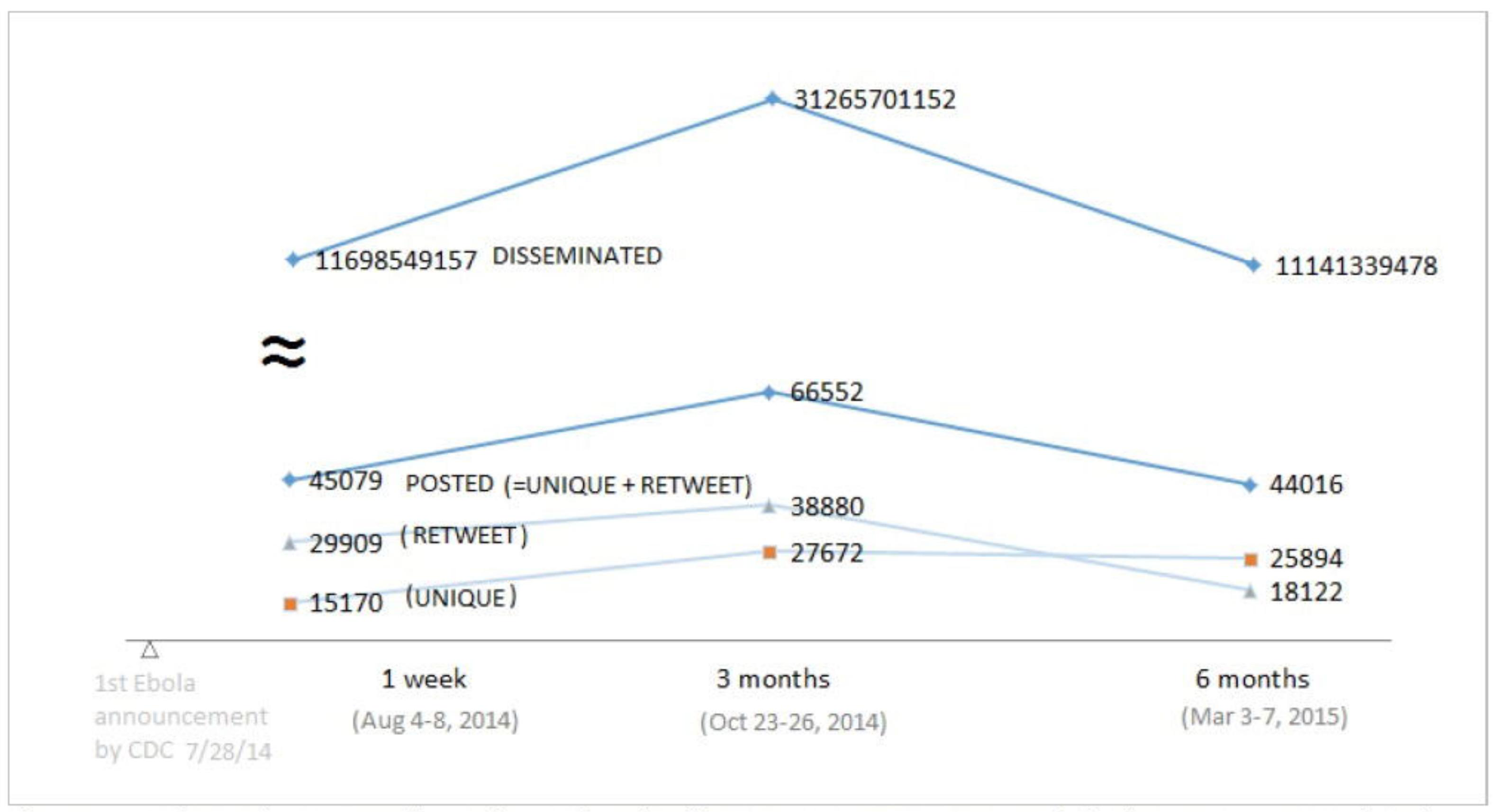
Information speed in Twitter after the first CDC announcement of Ebola (Tweets posted and disseminated to Twitter followers).

#### Information Seeking and Needs

Tweet topics at Time point 1 were: 1) Readiness of U. S. hospitals; 2) History and symptoms to report (e.g., travel to West Africa with high fever); 3) Ebola Statistics (i.e., number of death, number of cases, affected countries) and 4) Transmission mode (i.e., information about how Ebola spreads). Topics at Time point 2 were: 1) Response of Spanish government to Ebola (e.g., public frustration with its corrupted process of Ebola treatment); 2) Non-scientific information-seeking behavior of Ebola cure (e.g., magic healing of crystal stone for Ebola); 3) U. S. government policy (i.e., public response to the 21-day Ebola quarantine for volunteers from West Africa in New York and New Jersey); and 4) Fear of volunteering in West Africa (i.e., discouragement of volunteering for Ebola in West Africa such as Doctors Without Border). Topics at Time point 3 were: 1) Non-governmental organization (NGO) focusing on Save the Children and Mexico (e.g., protecting children in vulnerable Mexico); 2) Priorities and public frustration (e.g., 30 countries being depicted as vulnerable for Ebola and consequent public frustration towards slow response); 3) Lawsuit towards a hospital (i.e., Nina Pham, who was cured of Ebola, is suing the Texas Health Presbyterian Hospital in Dallas for lack of training and proper equipment); and 4) North Korean lifting of 4-month travel ban (i.e., public response to this), Table 1.

**Table I.**
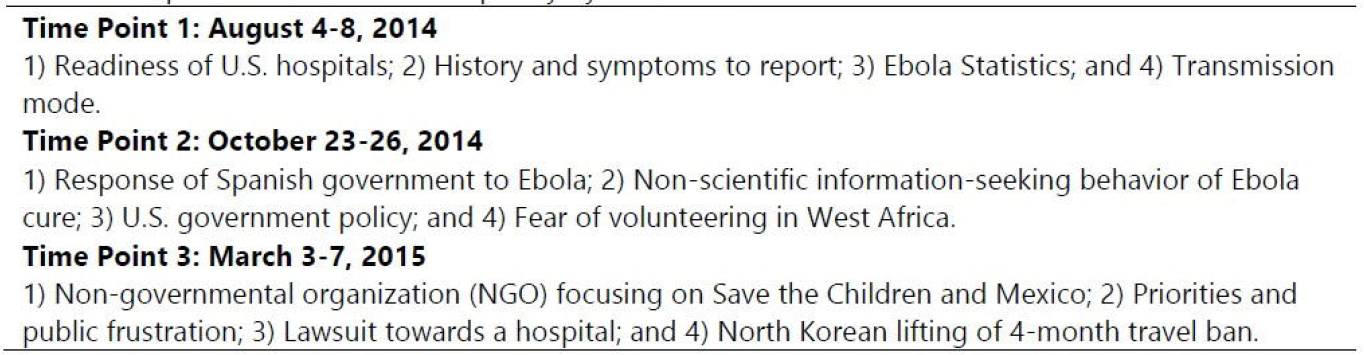
Top Tweets in Order of Frequency by Time Point

Main topics of concern discussed in tweets mentioning Ebola are illustrated with infographics in **Figure 2.** The bubble chart in Figure 2 illustrates the topics of Ebola tweets at each time point. The sizes of the bubbles were normalized by clusters representing their relative frequencies. The frequency of N-gram of the main topic for each time point is reported in Figure 2.

**Figure 2.**
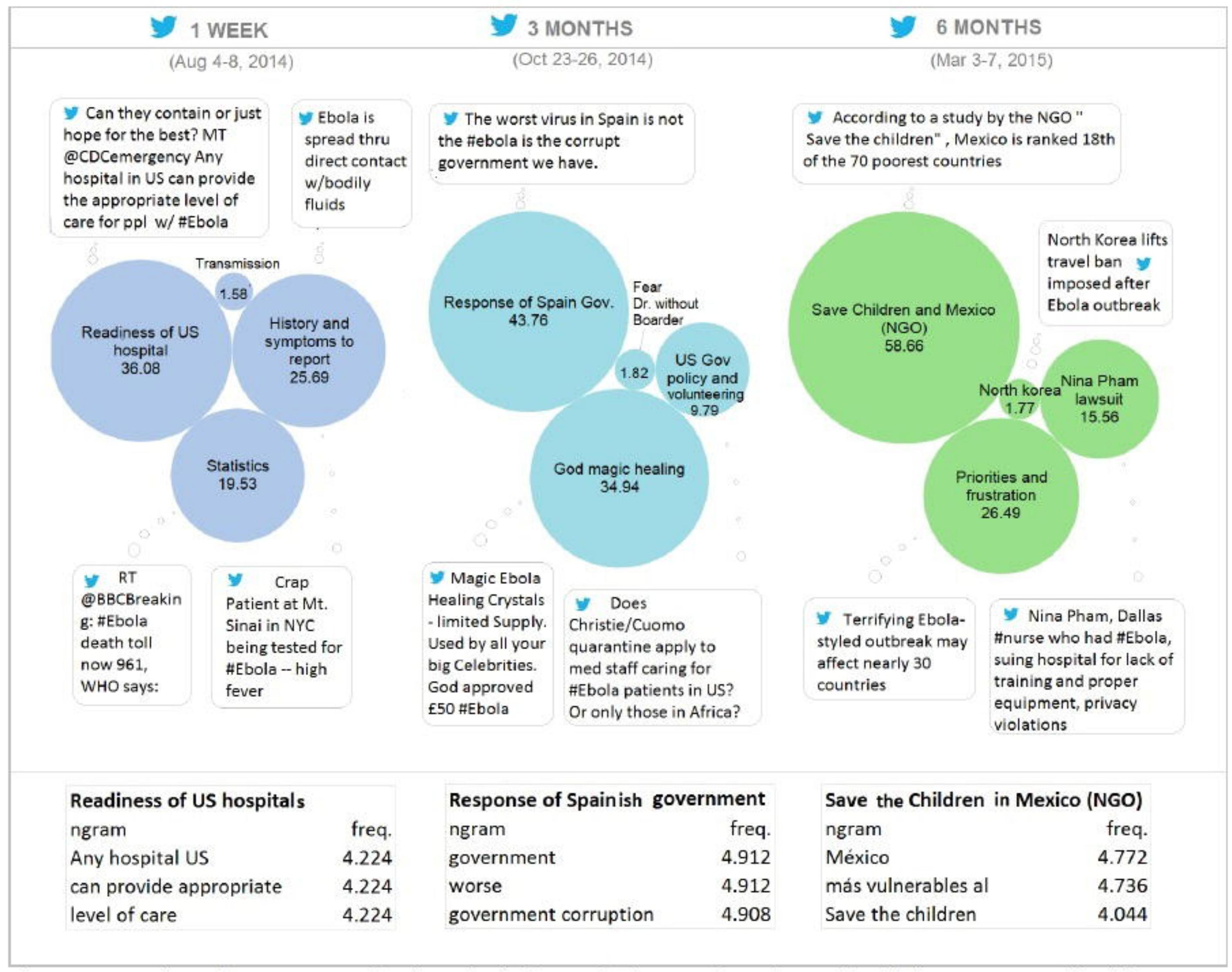
Topics of Tweets mentioning Ebola for each time point. Sizes of bubbles are normalized by clusters and show frequency of the topics.

#### In depth six-month analysis on News Announcements (March 3-7, 2015)

During March of 2015, as the epidemic came under control, information needs tweets declined. Results indicating Tweets by world region are displayed in Figure 3.

**Figure 3.**
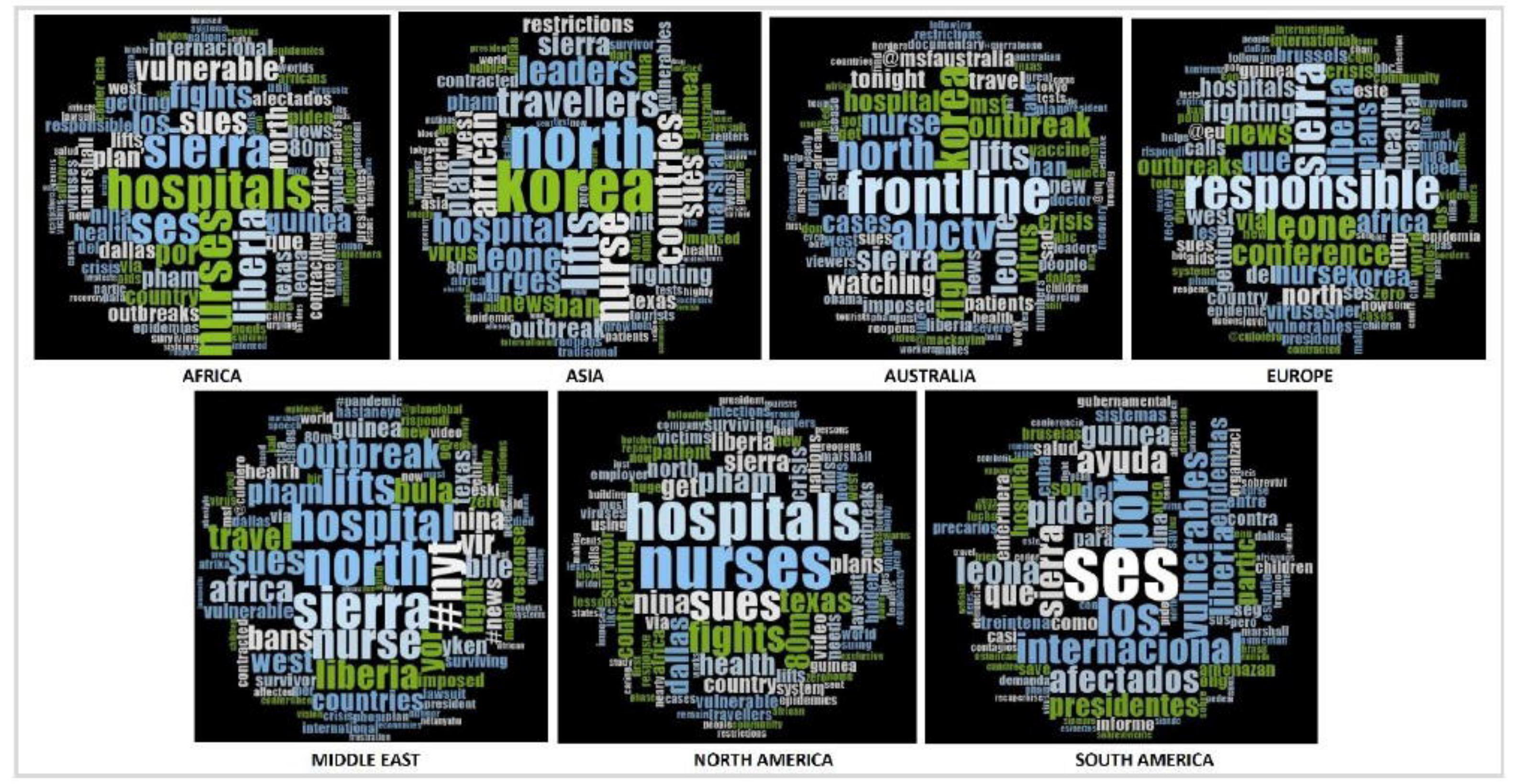
Ebola-related Tweets by world Region for the week of March 3-7, 2015.

During this period, tweets were regarding potential vulnerabilities to the virus, the Dallas Nurse lawsuit and the lifting of the North Korean travel ban. The one major information need still tweeted was Ebola-related vulnerabilities, all regions except Australia still felt vulnerable to Ebola exposure. In North America, Africa, Europe, Asia and the Middle East, top tweets focused around the Dallas nurse suing the hospital for Ebola exposure, Table 2.

**Table 2.**
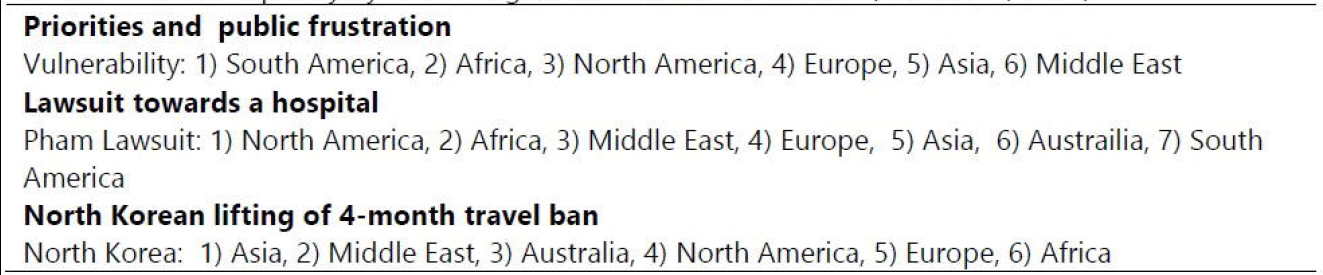
Top Frequency by World Region for 6 month Time Point (March 3-7, 2015)

Top Tweets were then ordered based on frequence and stratified by world region. Most frequent and consistent tweets across regions were Nurses (North America, Europe, Asia, Middle East and Africa); Hospitals (North America, Asia, Middle East and Africa); Sues (North America, Europe, Australia, Middle East and Africa) and Fight (North America, Europe, Australia and Africa), Table 3.

**Table 3.**
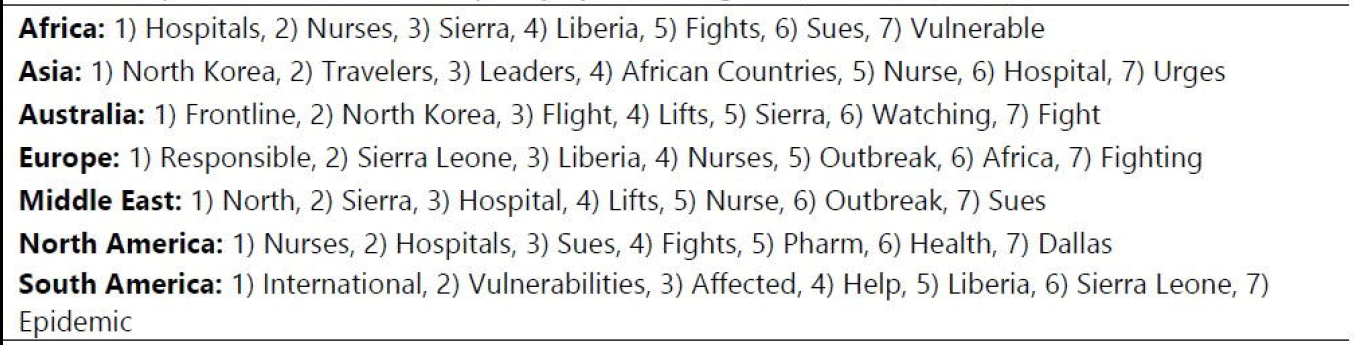
Top Tweets in Order of Frequency by World Regions (March 3-7, 2015)

## Discussion

Strategies for public health practice during outbreaks of emerging or reemerging infectious disease require a balance between protecting the health of the population and proper health education to minimize fear, discrimination and stigma^3, 11^. These difficulties mark the 2014-2016 Ebola outbreak. As the global community provided support to affected countries for containment and treatment of disease, fear spread unchecked throughout communities around the globe. Fear was driven by anxiety from a poor understanding of Ebola’s cause, infectivity and mortality^11^. The press further drove anxiety and fear as they reported worst-case scenarios and the possibility of a pandemic. It is evidenced that during outbreaks of epidemic or pandemic proportions, the public requires immediate health information^12^. The Conceptual Framework of PHSA demonstrates how health systems can link outbreak surveillance to action through data information messages^5^. However, the current study revealed through our content analysis of tweets at three time points during the 2014-2016 Ebola outbreak, information needs that persisted throughout the global community.

The foundation of outbreak surveillance is the delivery of useful and effective information whether to health officials to support disease containment or to develop tailored messages in health education campaigns^6, 13^. According to the Centers for Disease Control and Prevention (CDC), public health education efforts during outbreaks should include the surveying of attitudes and knowledge to support effective mass media messages^12^. Twitter and other social networking sites have the ability to support these education goals. These sites can be used as tools to both assess information needs and to disseminate tailored health information to address such needs. Our study, through content analysis, utilized Twitter to assess health information needs. In fact, tweets about the Ebola outbreak were posted in the tens of thousands with dissemination in the billions for all three-time points.

The week of August 4-8, 2014, reflects the early stage of health alerts when multiple countries confirmed Ebola cases and the fear of a global spread was at it’s peak. During this time point, Twitter users sought and disseminated Ebola information regarding concerns, transmission, infection and prevention. Top tweets were understandably regarding the readiness of US hospitals for potential cases. Concerns were followed by information needs, with the most frequent being the history of Ebola, symptoms, statistics on the epidemic and mode of transmission. This was also the time the first healthcare worker returned to the US (July 31, 2014)^11, 14^. Although the health worker was contained, Americans feared transmission and spread, indicating the tremendous need for health education campaigns. During this time period organizations such as Harvard and Columbia tweeted Ebola education messages, but dissemination was limited due to limited following, with no retweets from media sources.^18^

In the week of October 23-26, 2014, non-scientific information-seeking behavior of Ebola prevention and cure (e.g., magic healing of crystal stone for Ebola) was the second major topic tweeted and retweeted. During this period, Tweets discussed “God's magic healing crystal.” These findings clearly indicate health education campaigns were suboptimal in meeting the health information needs of the public. The end of October marked the first 3 months of extensive news coverage on the outbreak, travel bans and returning health professionals. Understandably, the second major topic tweeted was regarding the U. S. government policy surrounding containment, control and fear of those volunteering in West Africa. This was the ideal time to reinforce the modes of transmission, understanding signs and symptoms and plans for outbreak containment in the event of returning cases. Widespread reports from health entities including the CDC and the World Health Organization (WHO) also occurred early in the epidemic. In August of 2014, the WHO utilized Twitter to dispel social media rumors about such products claiming to cure and prevent Ebola^15^. However, three months later we can see that tweets about such remedies were persistent and concerns of outbreak containment remained apparent. These results revealed that information needs were not effectively met.

In the week of March 3-7, 2015, priorities and public frustration was the third major topic tweeted. This topic consisted of reports from non-governmental organizations (NGOs) about the list of 30 vulnerable Ebola countries. During this March time point, no further knowledge distribution regarding the spread and treatment of Ebola was observed as we saw in the beginning of August and the end of October. Tweets did continue around government policy for containment and control, specific to the North Korean lifting of the travel ban and the consequences of ineffective US hospital readiness with the their top tweets regarding the Dallas hospital lawsuit. Tweets during the March time point included ‘Ebola is not a disease that kills the rich in the West’. Such comments were retweeted by thousands. These tweets further indicate severe knowledge deficit. In fact, the 2014-2016 outbreak is the first in an urban setting with overcrowding driving the epidemic^3^. We cannot safely conclude that exposure to an infected person during rush hour on public transportation in a western city could not cause a problem. It is evident the need still existed for health information and effective education on outbreak containment and control.

An in-depth analysis of tweets the week of March 3-7 indicated that one major information need was vulnerabilities. All regions except Australia still felt vulnerable to Ebola exposure. In fact, South America, Africa and North America topped the list for frequent tweets about being vulnerable to Ebola. This is an indication of the continual and ongoing need for international health organizations to: 1) reiterate Ebola containment and control procedures and 2) to report the success of containment efforts and the effectiveness of airport and immigration screenings and quarantine. Such communication would have supported further reassurance of the public and remind them of their decreased vulnerability to Ebola exposure. Furthermore, this was an excellent time to reinforce the modes of Ebola transmission to diminish ongoing perceived vulnerabilities and fears. South America’s vulnerability concern stemmed from its close proximity to North America and Africa, tourism to their countries and the potential difficulty in managing or containing an Ebola outbreak^16^. This was an optimal time to reinforce educational messages, outbreak control and containment, to pledge support from developed nations and international organizations. Additionally, this was the ideal time to reemphasize appropriate screening protocols and to communicate successful containment efforts and approaches, particularly for developing countries such as Nigeria^18^.

In North America, Africa, Europe, Asia and the Middle East, top tweets focused around the Dallas nurse suing the hospital for Ebola exposure. This is direct insight into public sentiment about the actions taken by the nurse to sue for reckless endangerment. However, these messages are two–fold. They reinforced fear of deficiencies in the screening process, which made it possible for Mr. Duncan to leave the airport after arriving in Dallas. It also reinforced fears of hospital negligence. First, sending Mr. Duncan home and in turn exposing nurse Pham to Mr. Duncan’s most severe symptoms ^17^. This speaks to public sentiment regarding feelings of being unsafe. This again was a teachable moment allowing for the dispelling of myths, explaining what went wrong at the airport screening and at the Dallas hospital and discussing lessons learned. Information dissemination based on these tweets should have included new measures implemented to ensure improved screening procedures and protection for healthcare workers ^18^.

### Limitations

The generalizability of this study is limited due to the analysis of data in only two languages (English and Spanish) and a single social medium (Twitter). Our search strategies utilized all Ebola-related hashtag variations in an attempt to include the maximum number of Ebola-related tweets, with no analysis of their independent contributions to the results. Our tweet analysis focused on tweets developed or retweeted and not tweets that were only composed. Future studies can further support the understating of Ebola health information needs by exploring different social media in addition to blogs or community websites.

## CONCLUSION

Social media content can support outbreak surveillance efforts. Such sites allows for the development of networks to share and seek a variety of information regarding events including outbreaks such as Ebola. At the same time, social media can be an impediment to health security when it propagates (social contagion of simplistic ideas) information based on collective unawareness (e.g., crystals protect against Ebola). This phenomenon of social media fills a collective thought space that could be replaced with a wealth of scientific knowledge. Twitter and other social networking sites are also important for assessing health information needs and serving as platforms for information dissemination. Our study demonstrates how Twitter can support public health outbreak surveillance to identify health information needs. In our assessment, we have provided a window into public interests at three time points during the 2014-2016 Ebola epidemic. Ebola-related tweets were a rich source of experiences and opinions. Although a variety of health education efforts occurred during the outbreak, our results indicate health information needs were ongoing and transforming. Twitter can used as a knowledge transfer tool to assess the ongoing needs and support relevant data information messages during outbreak control and surveillance efforts. Content analysis of tweets provides an effective assessment of behavioral response and can in turn, support literacy-appropriate health information to improve behavioral response and diminish fear.

## Data Reporting

Data is owned by a third party, Twitter. The dataset of this study is found on a secure server please contact the author Dr. Sunmoo Yoon for access to the dataset.

